# User-friendly electron microscopy protocols for the visualization of biological macromolecular complexes in three dimensions: Visualization of *planta* clathrin-coated vesicles at ultrastructural resolution

**DOI:** 10.1101/2022.05.24.493253

**Authors:** Alexander Johnson, Walter A Kaufmann, Christoph Sommer, Tommaso Costanzo, Dana A Dahhan, Sebastian Y Bednarek, Jiří Friml

## Abstract

Biological systems are the sum of their dynamic 3-dimensional (3D) parts. Therefore, it is critical to study biological structures in 3D and at high resolutions to gain insights into their physiological functions. Electron microscopy of metal replicas of unroofed cells and isolated organelles has been a key technique to visualize intracellular structures at nanometer resolution. However, many of these protocols require specialized equipment and personnel to complete them. Here we present novel accessible protocols to analyze biological structures in unroofed cells and biochemically isolated organelles in 3D and at nanometer resolutions, focusing on Arabidopsis clathrin-coated vesicles (CCVs) - an essential trafficking organelle lacking detailed structural characterization due to their low preservation in classical electron microscopy techniques. First, we establish a protocol to visualize CCVs in unroofed cells using scanning-transmission electron microscopy (STEM) tomography, providing sufficient resolution to define the clathrin coat arrangements. Critically, the samples are prepared directly on electron microscopy grids, removing the requirement to use extremely corrosive acids, thereby enabling the use of this protocol in any electron microscopy lab. Secondly, we demonstrate this standardized sample preparation allows the direct comparison of isolated CCV samples with those visualized in cells. Finally, to facilitate the high-throughput and robust screening of metal replicated samples, we provide a deep learning analysis workflow to screen the ‘pseudo 3D’ morphology of CCVs imaged with 2D modalities. Overall, we present accessible ways to examine the 3D structure of biological samples and provide novel insights into the structure of plant CCVs.

## Main Text

Cellular processes are reliant upon the assembly and arrangement of organelles and macromolecular complexes. By defining the three-dimensional (3D) structure of these organelles and complexes we can gain critical insights into their physiological mechanisms and functions. This high-resolution 3D imaging of biological samples with nanometer resolution is routinely achieved by combining electron microscopy (EM) with tomographic acquisition protocols (1). An elegant and extremely useful methodology to prepare biological samples suitable for EM tomography is the metal replication of unroofed cells. This allows the direct visualization of the intracellular landscape and sub-cellular organelles *in vivo* (2-6). However, to examine these samples at nanometer resolutions via transmission electron microscopy (TEM), the samples must be on an EM grid. As many of the sample preparation protocols used for EM tomographic analysis of unroofed cells rely upon initially plating the cells onto glass coverslips, the resulting metal replicas must be transferred to EM grids by dissolving the glass with corrosive acids (4, 5); a procedure which must be conducted by facilities with specialized equipment and highly trained personnel, which restricts their wider use and application.

Clathrin-coated vesicles (CCVs) in mammalian cells are a good example of a well characterized organelle where their structures have been defined using EM methodologies for over 50 years (7). Consequently, we have a good understanding of their structural details and physiological functions in mediating cellular trafficking through the encapsulation of cargo in spherical membrane vesicles coated by a clathrin lattice (7-10). This clathrin lattice, composed of repeating clathrin triskelia formed of clathrin heavy and light chains, is arranged into pentagon and hexagon panels (resembling a honeycomb pattern) to create a 3D spherical coat/cage covering the vesicle (11-13). These CCVs are essential trafficking organelles formed during clathrin-mediated endocytosis, where they mediate cargo entry into the cell, and post Golgi-trafficking, to regulate protein sorting (8, 14-16). In stark contrast, while CCVs are also essential in plants (16, 17), we know very little about their structural and functional details. A major reason for this is because many of the standard EM sample preparation methods established in other model systems fail to reliably preserve *planta* CCVs. Furthermore, the TEM images of plant CCVs have been limited to planar two-dimensional (2D) views (17-19). Therefore, due to a lack of suitable protocols, a robust examination of the 3D structures of plant CCVs has been lacking.

To resolve the reliance on protocols which require specialist handling of acids, and facilitate the 3D analysis of plant CCVs at high resolution, we set out to establish protocols accessible to any routine EM lab. Using Arabidopsis CCVs as a model organelle, we describe an accessible novel metal replication protocol which can be performed directly on an EM grid; thereby removing the requirement for specialist equipment and personnel to handle extremely corrosive and hazardous reagents and enabling more researchers to visualize biological samples in 3D at high resolutions. We demonstrated that this standardized workflow produces 3D tomographic representations of single CCVs and allows the direct comparison of CCVs in unroofed cells with biochemically isolated CCVs, finding several subtle structural differences between endocytic CCVs with those biochemically isolated. In addition, we developed a semi-automated deep learning-based workflow which allows the robust high throughput 3D morphological examination of CCVs and validate this approach by examining CCVs structures in cells subjected to disruption of the endocytosis membrane bending machinery.

## Results

### Establishing scanning-transmission electron microscopy (STEM) tomography of CCVs in unroofed protoplasts derived from Arabidopsis seedlings (Protocol 1)

The unreliable preservation of plant CCVs *in situ* has hampered the characterization of these structure by EM methodologies. However, to overcome this we recently established a protoplast metal replica unroofing assay that reliably preserves large numbers of CCVs in plant cells (6, 20). Here, cells were attached to a glass coverslip, ‘unroofed’ (physically disrupted) to remove any cellular materials not associated with the plasma membrane (PM) directly attached to the coverslip, fixed, and coated with platinum; thus producing a metal replica of the inside cellular environment of plant cells which can then be examined with EM. As this approach relies upon the use of glass coverslips, visualization of CCVs was limited to 2D views and lacked the resolution to clearly determine the arrangement of the clathrin coat lattice. While we were able to overcome these issues to produce tomographic visualizations of single CCVs (6), we relied upon a specialist nanofabrication facility to use hydrofluoric acid to transfer the replicas from glass to EM grids – which is not accessible for many research labs. Thus, to further advance our understanding of the ultra-structural details of plant CCVs, we developed a novel method to produce high resolution 3D tomographic reconstructions of CCVs in plant cells directly adhered onto EM grids, bypassing the requirement for corrosive acids to allow any EM capable laboratory to contribute to the collective knowledge of plant CCVs.

We focused our method on utilizing protoplast cells derived directly from Arabidopsis seedlings (Fig 1A). As the cells are prepared and processed directly upon EM grids, a critical step of the method relies upon the cells having a good attachment to the Formvar film on the EM grid. This is because the cells need to adhere strongly enough to remain attached during multiple washing steps and still have a section of the PM attached during the physical unroofing step. We chose to utilize gold EM grids with a Formvar film to remove the possibility of metal toxicity during the incubation steps of this protocol. Then we treated these EM grids with a carbon and poly-l-lysine coat to aid the attachment of the cells, which was achieved by letting the cells settle into the grid during a 3-hour incubation (Fig. S1). Once attached, the further steps were conducted on small droplets of solution on parafilm sheets due to their small surface area and the fragility of the EM grids. This also enables experimenters to process multiple samples with small volumes of reagents and materials.

**Fig. 1.**
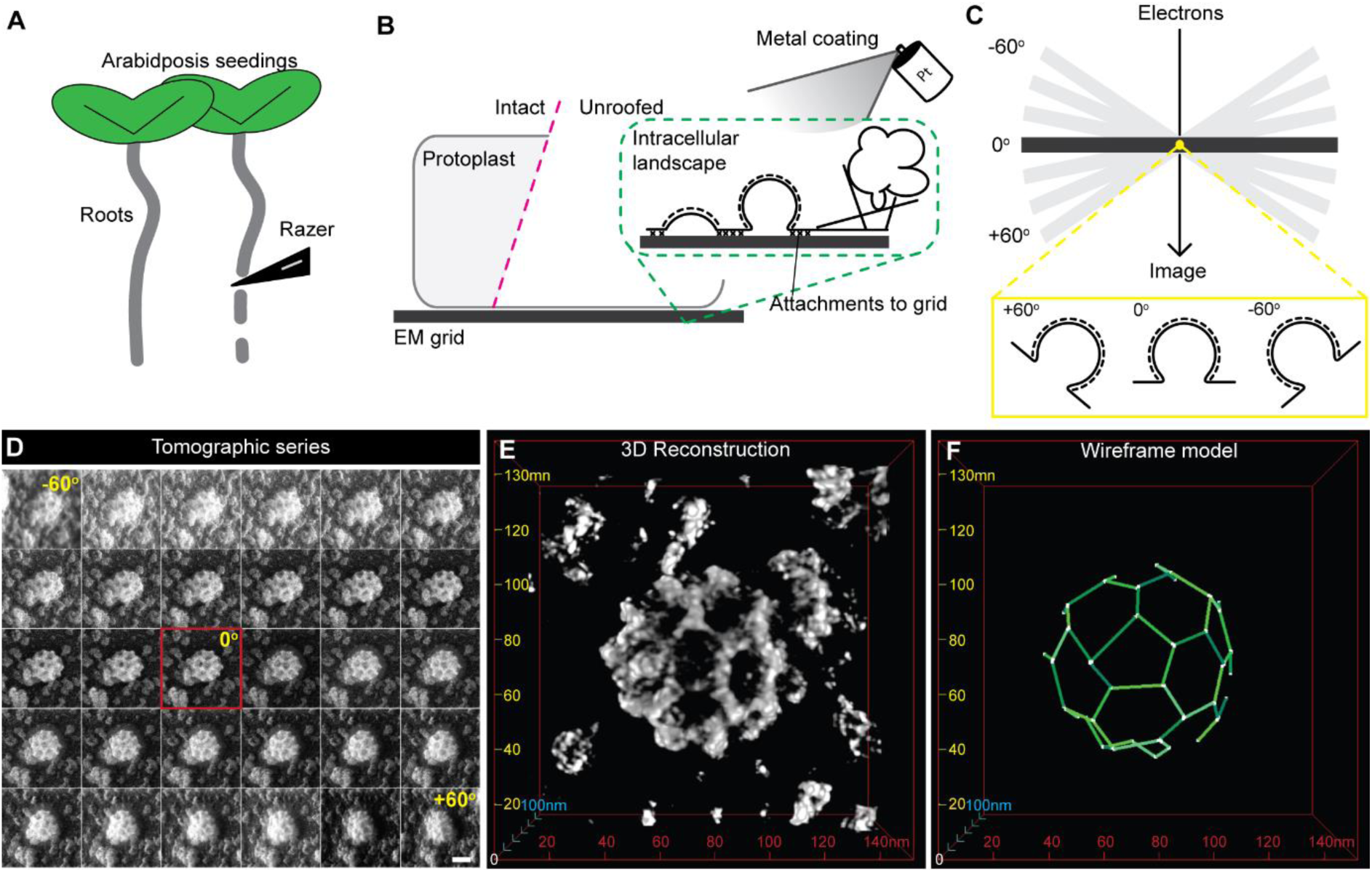
STEM tomography of CCVs in metal replicas of unroofed protoplasts derived from Arabidopsis seedlings. (A) Roots from Arabidopsis seedlings are dissected and treated to produce protoplast cells. (B) The cells are then attached directly to EM grids and unroofed (magenta dashed line) such that only the plasma membrane and its associated structures remain attached to the EM grid. This is then coated with metal to make a replica of the intracellular landscape. (C) The replica is visualized using EM tomography, where the sample is tilted through a series of angles allowing many perspectives of CCVs to be acquired (yellow dashed box). (D) Example STEM tomogram of a CCV in a metal replica of an unroofed Arabidopsis root protoplast. Scale bar, 50 nm. (E) A 3D reconstruction of ‘D’. (F) A wireframe model of the reconstructed tomogram is produced to enable quantification of the structural details of the CCV.

Given the fragility of the EM grids, it was not possible to unroof the cells with mechanical disruption methods like sonication, which is typically used in unroofing protocols (5). Therefore, we relied upon the application of a detergent (Triton X-100) with gentle agitation of the samples to disrupt the cellular membranes. In addition, to dehydrate the samples rapidly without risking damage to the EM sample grids, we utilized manual application of hexamethyldisilane rather than relying on a critical point dryer device (21). The samples were coated with platinum to produce a replica of the unroofed cell, and an additional layer of carbon was applied to enhance the stability of the sample and to prevent electron contamination during image acquisition (Fig. 1B).

To visualize the metal replicas of the unroofed cells, and capture 3D images of the CCVs, we employed STEM tomography. We focused exclusively on CCVs formed during clathrin-mediated endocytosis by examining only PM-associated CCVs. Tomograms of individual CCVs were generated by tilting the EM grid through a range of angles (typically -72° to +72°) along a single tilt axis and images from each perspective were acquired (Fig. 1C and D). The tomogram images were aligned by cross-correlation and used to create a 3D visualization of the CCV (Fig. 1E and Movie S1). From these 3D reconstructions we were able to precisely determine the structural details of the CCVs, such as diameters, volumes, arrangement of clathrin lattices and lengths of the triskelia arms, by making wireframe models of the CCVs (Fig. 1F and Table S1). As such, we found that the average Arabidopsis root epidermal cell endocytic CCV has a diameter of 75.99 ± 0.63 nm, volume of 1868.5 ± 45.54 μm^3^ and 17 clathrin panels (5 pentagons, 4 hexagons, and 8 not fully visibly closed at the base of the CCV) with a branch arm length of 16.88 ± 0.06 nm.

Thus, we established an accessible user-friendly ‘on-grid’ metal replication method, which removes the requirement to chemically dissolve glass coverslips, allowing the direct 3D STEM nanometer resolution imaging of CCVs in cells, thereby revealing details of the *planta* CCVs clathrin coat arrangements.

### 3D analysis of biochemically isolated CCVs prepared using the ‘on-grid’ protocol (Protocol 2)

To demonstrate the versatility of the ‘on-grid’ metal replication protocol, we examined the 3D ultrastructure of CCVs biochemically isolated from plant cells. Here, we employed the same protocol for preparing unroofed cells to prepare and visualize purified CCV preparations, allowing the direct comparison of isolated and *in vivo* organelles, such as purified CCVs with *bona fide* endocytic CCVs.

CCVs were isolated from undifferentiated Arabidopsis suspension cell cultures (22), and attached directly to glow discharged and carbon coated EM mesh-grids. The samples were then processed with the same protocol as unroofed cells. Imaging the metal replicas of purified CCVs with STEM tomography allowed the production of 3D reconstructions of the isolated CCVs (Fig. 2A, B and Movie S2), enabling their quantitative structural analysis (Fig. 2C). We found that the average isolated CCV was 71.06 ± 1.65 nm in diameter, 1547.45 ± 110.64 μm^3^ in volume and comprised of 17 clathrin panels (6 pentagons, 4 hexagons, and 7 not fully visibly closed at the base of the CCV) with a branching arm length of 18.22 ± 0.14 nm (Table S2).

**Fig. 2.**
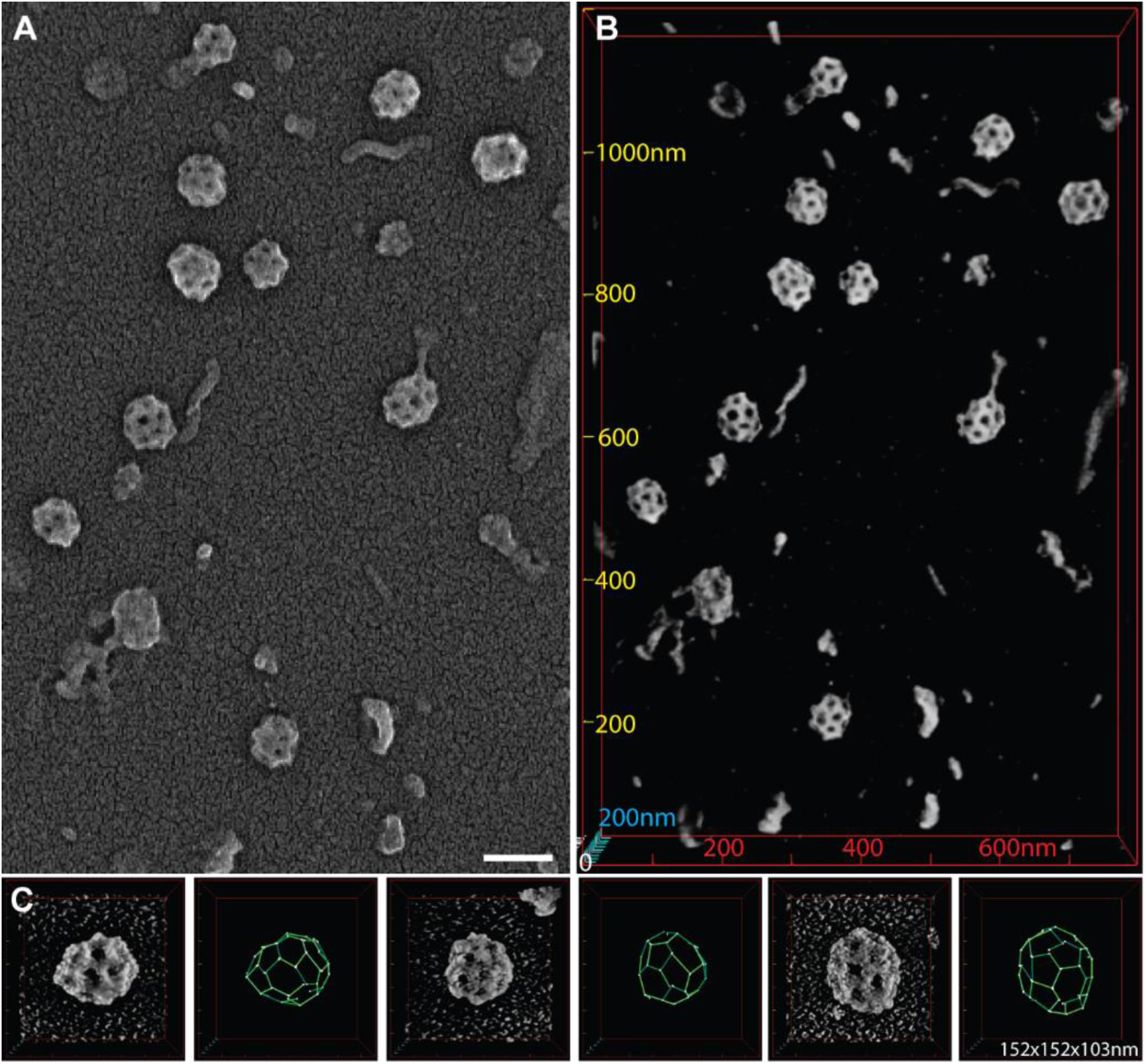
Application of the standardized ‘on grid’ metal replication to visualize biochemically isolated CCVs in 3D. (A) Example STEM image of metal replicas of isolated CCV preparations from Arabidopsis T87 suspension cell cultures. Scale bar, 100 nm. (B) A 3D reconstruction of ‘A’. Scale values are reported as nm. (C) Representative of 3D reconstructions and wireframe models of single CCVs. 3D volume; x = 152, y = 152 and z = 103 nm.

This demonstrates the versatility of the on-grid metal replication protocol and provides a standardized protocol to structurally characterize CCVs from different biological sources using STEM tomography.

### High throughput pseudo 3D morphology screening of CCVs (Protocol 3)

While tomographic reconstructions of single CCVs provides the direct visualization of their 3D structure and morphology at high resolution, it is a low throughput approach. Thus, to increase our ability to examine large numbers of CCV structures, we developed a deep learning-based workflow to analyze the morphology of CCVs from images acquired at lower resolutions and in a single plane. This morphologic analysis provides a rapid and robust ‘pseudo 3D’ analyses of CCVs and their formation in a high throughput and unbiased manner.

To develop this method, we used images of metal replicas of unroofed protoplasts derived from Arabidopsis root cells attached to glass coverslips (6). Here, as the metal replicas were imaged in 2D with a large field of view using scanning electron microscopy (SEM), each image contained many visible CCVs enabling the high throughput visualization of CCVs. To characterize the morphology of the CCVs, we quantify various object features: such as area, maximum and minimum diameters, and average grey value. To date, such analysis has relied upon manual segmentation of the CCVs by the experimenter. To increase the throughput and reproducibility of the morphology quantification of CCVs, we used deep learning to create an accurate CCV segmentation model. To do this, we created a comprehensive training set of image pairs; consisting of raw images together with these same images where the CCVs are manually annotated (Fig. 3A). This training dataset was then used to train a state-of-the-art neural network model called Cellpose to generate an accurate CCV prediction model (23). To increase the robustness and the ability of the model to predict all possible CCV structures, the training set included images of metal replicas of unroofed cells derived from wild-type plants and from a mutant that produces clathrin plaques (which are absent from wild type cells (6, 20, 24)). The trained model was then used to automatically detect CCVs in additional images which had not previously been seen by the model (‘unseen images’) (Fig. 3B). To validate the model predictions, we compared them to manual segmentations on a dataset of CCVs from wild-type cells. From 17 SEM images of metal replicas of unroofed cells, manual segmentation identified 234 CCVs, whereas the automated method successfully predicted 232 (Table S3). To determine if the CCV segmentations from our model matched those made manually, we calculated an average pixel overlap of 83% between the manual and automated CCV segmentations using an intersection over union (IoU) calculation (Fig. S2), indicating the Cellpose model predicted the same CCV structures and with high degree of accuracy. The predicted CCV segmentations were then exported as regions of interest (ROIs) to FIJI and quantified. This ‘FIJI analysis’ step was not automated to provide a checkpoint for users to visualize and confirm the automated CCV segmentations. We found that the average areas reported by manual and automated analyses were not significantly different (t-test, p = 0.35) (4.93 ± 0.21 μm^2^ (dimeter of 79.24 nm) for manual and 5.24 ± 0.25 μm^2^ (dimeter of 81.71 nm) for the automated segmentations (Table S3)), thus validating the accuracy of this approach.

**Fig. 3.**
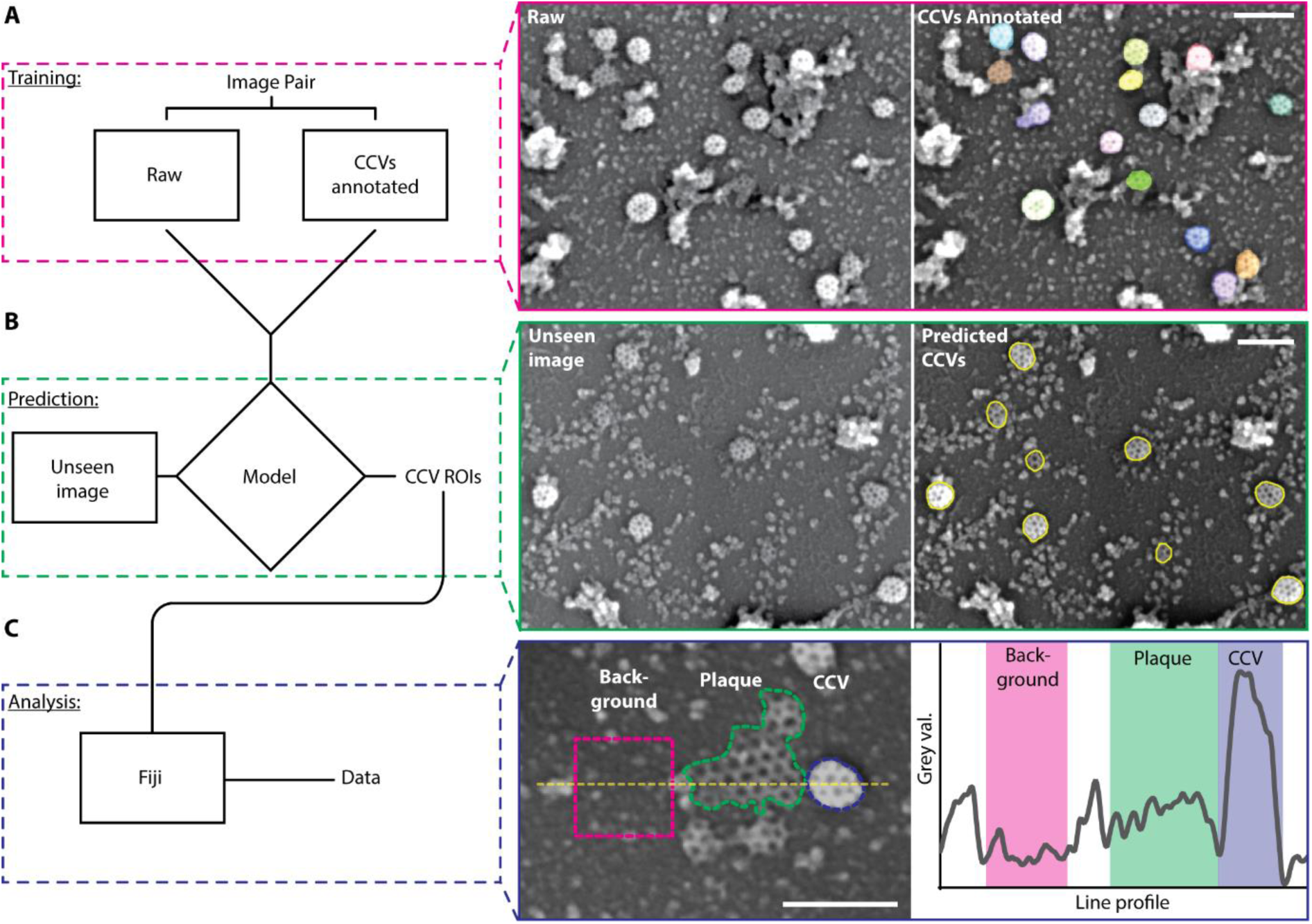
High throughput deep learning analysis of CCV pseudo-3D morphology. (A) Model training: example image from the training set with manual annotation (left hand image, multiple colors) overlaid of metal replicas of unroofed Arabidopsis root protoplasts acquired by SEM. (B) Model prediction: the model was used to predict CCVs in images not included in the training set (“unseen image”), which is outputted as a series of ROIs of CCVs (yellow outline). (C) The predicted CCV ROIs are opened in Fiji to quantify morphological features of each CCV. (Blue box) a pseudo 3D metric is provided by the gray value of the CCV as the more spherical a CCV is, the higher its grey value; the grey values of the PM (magenta dashed line), a clathrin plaque (green dashed line) and typical CCV (blue dashed line) are measured along the yellow dashed line and is quantified on the left plot. Scale bars, 200 nm.

To provide a pseudo 3D metric, CCV curvature, the average grey value of the CCV was used as a proxy for how spherical/3D the CCV is. During its development, the CCV becomes more protein dense and projects away from the PM, yielding a higher grey value than flat structures on the PM (4) (Fig. 3C). By normalizing the CCV grey values to an area of flat PM in each image, we can generate relative estimates of 3D curvature and compare them across different images. Comparing the curvature estimation from the manual and automated segmentations from the validation wild-type data resulted in similar values; 2.23 ± 0.07 arbitrary units (au) for manual and 2.19 ± 0.07 au for automated segmentations (Table S3).

To highlight the robustness of this protocol in identifying and screening the ‘pseudo 3D’ morphology of CCVs in a high throughput manner, we applied it to SEM images of unroofed cells where the clathrin-mediated endocytosis membrane bending machinery was disrupted (Fig. 4A). These images were of metal replicas of unroofed protoplasts derived from the roots of *WDXM2* seedlings, an inducible loss-of-function TPLATE mutant that prevents the formation of spherical CCVs (6, 25), subjected to control (non-induction) and disruptive (induced) conditions. The automated segmentation successfully detected the clathrin structures during both experimental conditions, finding one additional CCV compared to manual segmentation for each condition. The IoU pixel overlap was 92% when comparing the manual and automated segmentations (Table S4), indicating that the structures identified had a high degree of overlap. The area and the estimated 3D curvature values of each CCV were plotted against each other for each experimental condition (Fig. 4B), showing the 3D morphological differences of the CCVs in control and membrane bending disruptive conditions. To further define these morphological differences, we imposed thresholds upon the data (4). First, an area threshold of 8500 nm^2^ (diameter of ∼105 nm) was used to classify clathrin structures as ‘large’ or ‘small’. To classify clathrin structures based on their 3D shape, we identified ‘curved’ or ‘flat’ structures, which were defined by the average curvature estimation of the 3 flattest CCVs visible in the control conditions determined manually (1.54 au). Combining these thresholds created 4 morphological categories to define the clathrin structures: ‘small and round’ (SR) representing productive endocytic CCVs, ‘small and flat’ (SF), ‘large and round’ (LR) and ‘large and flat’ (LF) representing plaques. We found that the majority of CCVs formed during the control conditions are ‘small and round’ (94% compared to 13% during disruption), whereas the majority of CCVs formed during membrane bending disruption were found to be small and flat (70% compared to 4% in control) (Fig. 4B). This agrees with our previous results examining CCVs in metal replicas of unroofed *WDMX2* protoplasts (6), which were based upon time consuming manual segmentations and high-resolution STEM tomography therefore confirming the utility of the automated segmentations for high-throughput screening of the pseudo 3D morphology of CCVs.

**Fig. 4.**
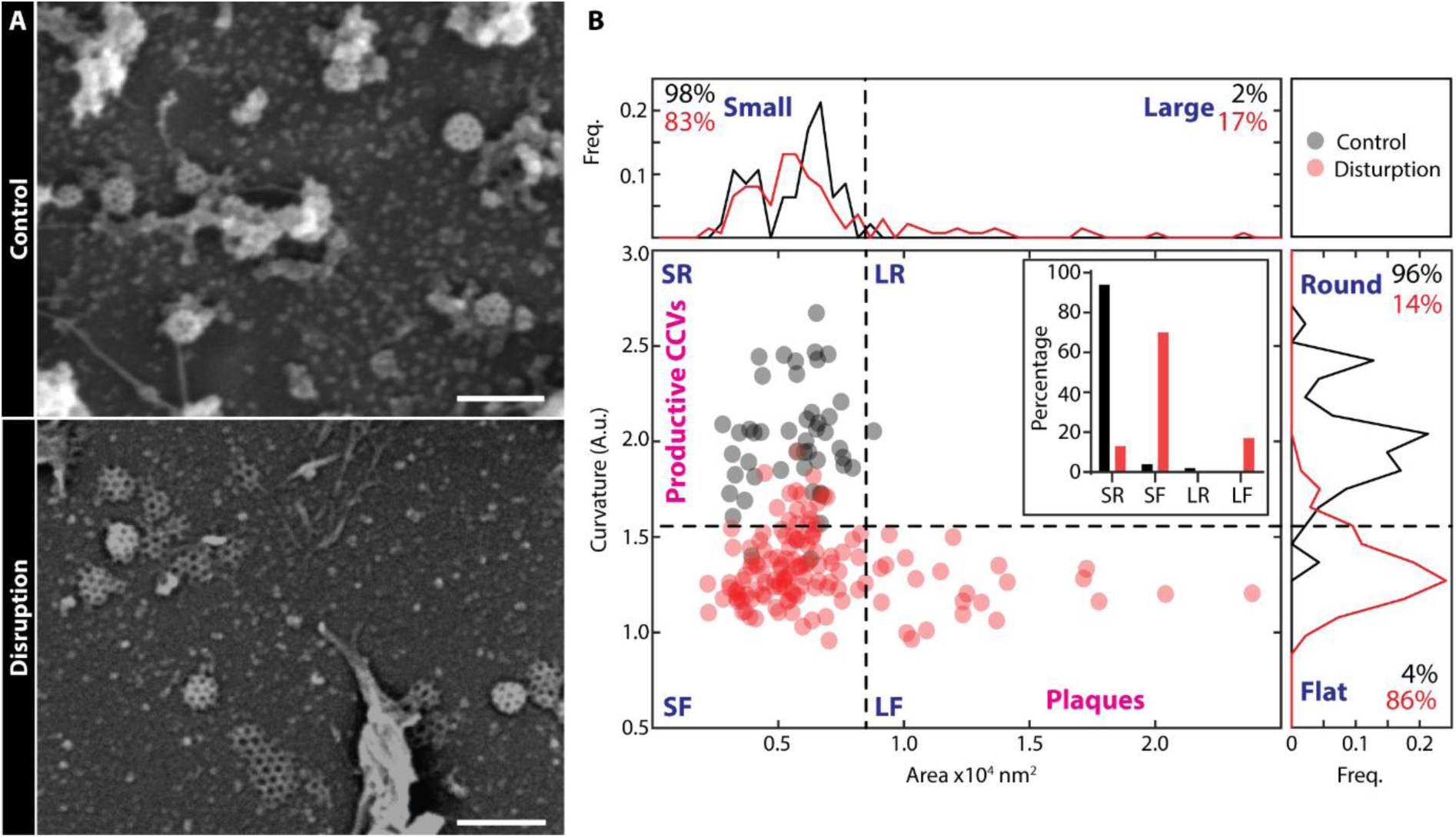
Deep learning analysis of CCV morphology during membrane bending disruption. (A) Example SEM images of metal replicas of unroofed protoplasts derived from roots of *WDXM2* seedlings subjected to control or membrane bending disruptive conditions. Scale bar, 200 nm. (B) Quantification of CCV morphology as detected using our deep learning-based analysis in control (black) or disruptive (red) conditions. Histograms (upper and right panels) and scatter plot (bottom left panel) of the areas and estimated 3D curvature of CCVs in control (black) or disruptive (red) conditions. Thresholds are used to classify the CCVs (black dashed lines) into categories based on their 3D shape; SR (small and round), SF (small and flat), LR (large and round), and LF (large and flat, plaques). The percentage of CCVs in each category is shown in the insert bar graph. N; control, 47 CCVs; disruption, 131 CCVs.

This, we provide a robust workflow which allows the high throughput pseudo 3D morphological screening of organelles in unroofed cells, enabling robust quantitative analysis of CCVs in cells subjected to physiological disruptions.

## Discussion

Understanding the structural details of biological samples provides key insights into their functions, therefore high-resolution 3D imaging is an extremely powerful tool for investigating cell biology. While EM techniques are routinely used to achieve this, they are traditionally often reliant upon harsh and/or dangerous chemicals requiring specialized facilities to conduct them (5, 6), which can limit their accessibility to the biological community. To overcome this, we developed protocols which allow the 3D examination of metal replicated samples in any routine EM lab (Fig. 5). Specifically, we established a method for preparing samples (unroofed cells and biochemically isolated organelles) directly on EM grids, providing a standardized sample preparation protocol which does not require the use of extremely toxic acids. These preparations can be combined with STEM tomography to produce nanometer resolution 3D images. However, as this high-resolution imaging can be time consuming, we also developed a high throughput deep learning-based analysis which provides a protocol for rapid morphological ‘pseudo 3D’ screening of biological structures. While these protocols can likely be adjusted for a range of cell types and organelles, here we applied them to further characterize the 3D morphology of plant CCVs - an essential trafficking organelles underpinning many cellular processes (26) - at high resolutions. We first visualized the clathrin assembly of CCVs in unroofed protoplasts to investigate endocytic CCV structures. By using the same sample preparation for biochemically isolated CCVs, we could directly compare them to specific CCV populations visualized inside cells. While we found that there were small differences in the sizes of endocytic and isolated CCVs, this is likely due to the fact that the isolated CCV preparations from whole cell lysates contain a mixture of CCVs derived from different origins - clathrin-mediated endocytosis and the early endosome/trans-Golgi network (22, 27, 28). We then validated our high throughput pseudo 3D analysis by applying it to metal replicas of unroofed cells carrying an inducible loss of endocytic membrane bending.

**Fig. 5.**
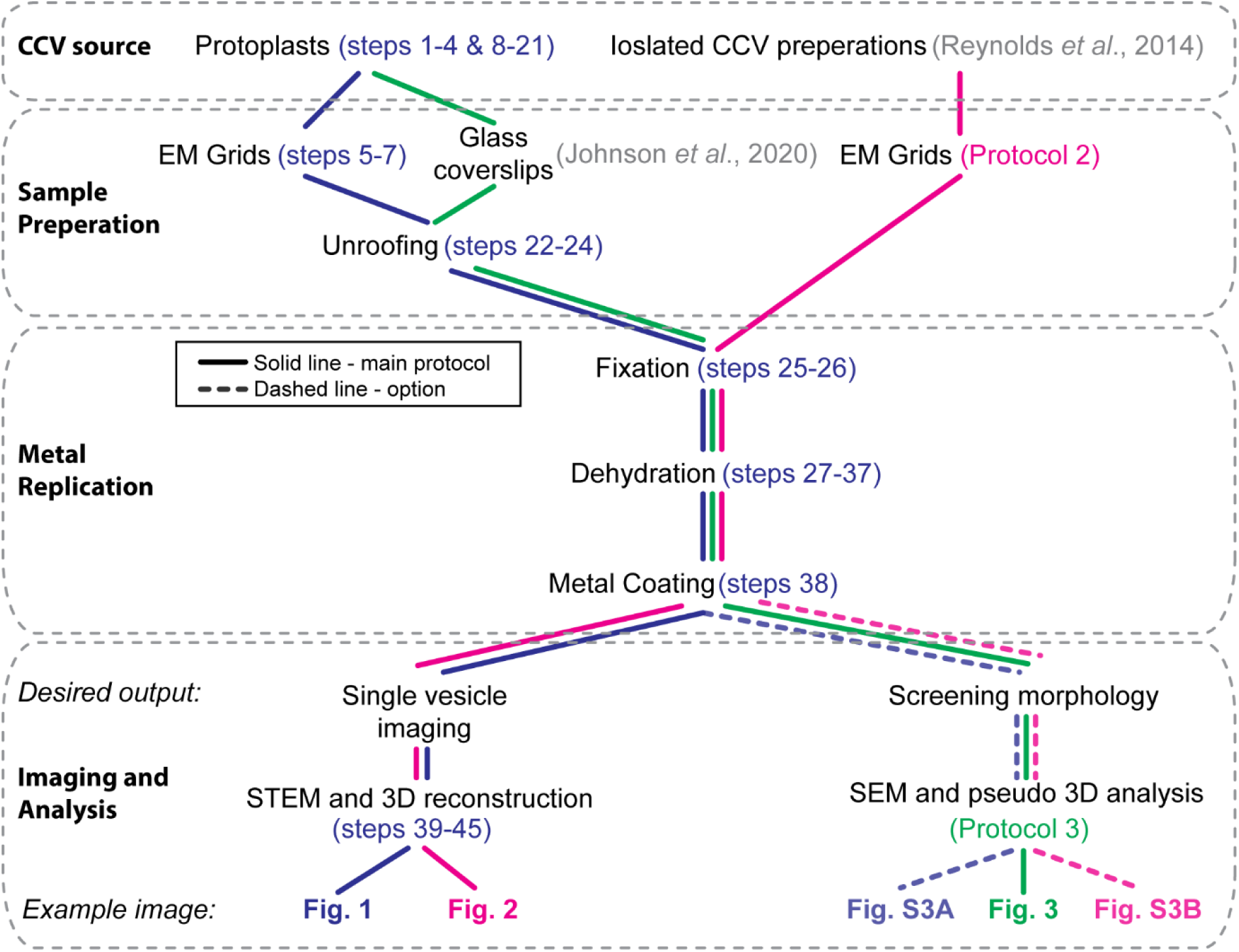
Overview of methods to examine plant CCVs in 3D as presented in this manuscript. Flow chart showing the versatility of the presented protocols to visualize CCVs in 3D. Steps refer to the detailed step-by-step protocols included in the ‘Supplemental Materials and Methods’ (blue; Protocol 1 - STEM Tomography of unroofed protoplast cells derived from Arabidopsis seedlings; magenta, Protocol 2 - STEM Tomography of isolated CCVs derived from cultured Arabidopsis cells; green, Protocol 3 - high throughput pseudo 3D morphological screening of CCVs).

To produce samples which reliably contain *planta* CCVs, we made use of protoplasts. This is because for an unknown reason, CCVs are rarely preserved no matter which EM sample preparation is used with whole plant samples. Therefore, to produce any robust quantitative analysis we had to utilize a cell system which allowed the consistent preservation of CCVs. As a result, we are examining plant CCVs which have formed in a different biophysical cellular environment compared to those in intact plants. However, protoplasts contain the same protein machinery responsible for producing CCVs, and the diameters of the protoplast CCVs (76 nm) are similar to the few reported measurements made in intact plant tissues (∼80 nm (17, 18)), thus highlighting that the difference in cellular environments does not affect the overall production of CCVs.

We focused on utilizing protoplasts derived directly from Arabidopsis seedings. This allows one to use already existing and established genetically altered plants as starting material. Furthermore, it provides the opportunity to reliably examine the effect of mutants upon CCV formation and structure. The use of homozygous Arabidopsis seedlings ensures that every cell unroofed carries the genetic manipulation, which is not possible to guarantee using transient transfection protocols which are not 100% efficient.

Our high-throughput screening of CCVs pseudo 3D morphology analysis was developed using samples attached to glass coverslips. This demonstrates that 3D analysis of samples can be conducted even without high end STEM enabled microscopes. To test the versatility of this analysis we applied the same CCV segmentation model to SEM images of metal replicas of unroofed protoplasts prepared directly on EM grids (Protocol 1) and to isolated CCVs (Protocol 2) (Fig. S3). We found that the model was accurate for both the in cell and purified CCVs. However, to further improve this accuracy, we suggest training specialized models using representative CCV images of the specific preparation method of interest.

We also expect our protocols to become a platform for further technical innovations and optimizations to continue advancing our understanding of these critical trafficking organelles. For example, combining these methods with immunolabeling approaches and fluorescence super resolution microscopy, such as correlated light and electron microscopy (CLEM) procedures, would help to unravel the molecular composition of CCVs and allow the precise localization of CCV related proteins. These sample preparation methods could also aid researchers to further develop focused ion beam (FIB) milling approaches to capture images of CCV *in situ* inside plant tissues.

Overall, we present refined accessible user-friendly methods for the 3D analysis of biological samples and investigate the 3D structural details of plant CCVs at unprecedented resolutions and accuracy. The experimental workflows provide the opportunity for the direct comparison of CCVs derived from differing sources, like specific cell types and purified CCV preparations. Additionally, the methods allow one to define individual structures at high resolution and screen CCVs in a high throughput machine learning assisted analysis to robustly quantify their morphology.

## Materials and Methods

For detailed step-by-step protocols of the methodologies presented in this manuscript (Protocol 1, STEM tomography of unroofed protoplast cells derived from Arabidopsis seedlings; Protocol 2, STEM tomography of isolated CCVs derived from cultured Arabidopsis cells and; Protocol 3, high throughput pseudo 3D morphological screening of CCVs), please refer to the Supplemental Materials and Method section.

### Plant materials and sample preparation

To generate Arabidopsis root protoplasts, Arabidopsis Col-0 seeds were sown on ½ AM agar plates, supplemented with 1% (w/v) sucrose. The plates were stratified by incubation at 4°C for 2-3 days in the dark and then grown vertically for 8-10 days (21°C; 16 hours light and 8 hours dark cycles). The roots were dissected from the seedlings and then cut into small sections (∼1-2 mm) in ‘enzyme solution’ (0.4 M mannitol, 20 mM KCl, 20 mM MES pH 5.7, 1.5% cellulase R10 [Duchefa #C8001], and 0.4% macerozyme R10 [Serva #28302] in H_2_0). The enzyme solution and sections were then incubated in a vacuum chamber for 20 minutes and then incubated at room temperature and atmospheric pressure in the dark with gentle agitation for 3 hours. The cells were collected by centrifugation at 100 rcf (relative centrifugal force) for 2 minutes at room temperature. The pelleted cells were washed with W5 buffer (154 mM NaCl, 125 mM CaCl2, 5 mM KCl, and 2 mM MES) by centrifugation (100 rcf for 2 minutes). The cells were resuspended in W5 buffer and incubated at 4°C for 30 minutes. The cells were then centrifuged (100 rcf for 2 minutes) and resuspended in ‘hyperosmotic growth media buffer (0.44% [w/v] MS powder with vitamins [Duchefa #M0222], 89 mM sucrose, and 75 mM mannitol, pH5.5 adjusted with KOH) and plated on to pre-prepared EM grids (detailed in the following section), where they were incubated in a humid chamber for 3 hours. The cells were then washed with PBS. To unroof the cells, they were then incubated with extraction buffer (2 μM phalloidin, 2 μM taxol, 1% (w/v) Triton X-100 and 1% (w/v) polyethylene glycol (PEG; MW 2,000) in PEM buffer (100 mM PIPES free acid, 1 mM MgCl2, 1 mM EGTA; pH 6.9 adjusted with KOH) for 5 minutes with gentle agitation. The samples were then washed three times with PEM buffer supplemented with 1% (w/v) PEG 2,000. Samples were fixed using 2% (w/v) glutaraldehyde (GA) in 0.1 M phosphate buffer (PB) via a 30-minute incubation. The samples were then washed twice with 0.1 M PB and stored in 0.1 M PB at 4°C until further processing.

Purified CCVs samples were isolated from undifferentiated T87 Arabidopsis suspension cell cultures by differential gradient centrifugation, as previously described (22). 5 μl of the CCV preparation ([0.33 mg/ml]) was plated and incubated on a pre-prepared EM grid (detailed in the following section) for minutes. Whatman™ blotting paper was used to remove the excess solution. The samples were then fixed by a 30-minute incubation with 2% GA in 0.1 M PB via a 30-minute incubation.

### EM grid preparations

Gold EM grids with Formvar-film (Electron Microscopy Sciences #G300PB-Au) were used for processing protoplast samples. They were coated with carbon to a thickness of 10 nm using a Leica ACE600 coating device. The grids were then supplemented with poly-l-lysine (Sigma) by a 15-minute incubation and washed with H_2_O.

Copper EM grids with carbon-film (Electron Microscopy Sciences #CF300-Cu) were used for isolated CCV preparations. They were subjected to glow discharge (4 min at 7×10E-1 mbar) using the ELMO glow discharge system (Agar Scientific Ltd.).

### Metal replication of samples

The fixed samples were washed with 0.1 M PB and H_2_O, and then incubated with 0.1% (w/v) tannic acid for 20 minutes. The samples were washed three times with H_2_O and incubated with 0.2% (w/v) uranyl acetate for 20 minutes. To dehydrate the samples, they were washed three times with H_2_O, infiltrated with graded ethanol (10%, 20%, 40%, 60%, 80%, 96% and 100%) and subjected to a 2-minute incubation with hexamethyldisilane. The samples were then dried by evaporation. The samples were then coated with 3 nm platinum and 4 nm carbon using the ACE600 coating device (Leica Microsystems).

### STEM tomography

STEM tomograms were acquired with a JEOL JEM2800 scanning-transmission electron microscope (200 kV AV) control by STEM Recorder (https://temography.com/en/). Diameters of the CCVs were determined from the 0° single plane STEM image, where an average value was calculated from the maximum and minimum Feret distances of a ROI drawn around the CCV in Fiji (29). This value was then used to calculate a spherical volume. Tomograms of single CCVs were recorded over a range of ∼-72° to +72° along single tilt axis with step size 4°. They were aligned and reconstructed using Composer (https://temography.com/en/). Wireframe models of the CCVs were manually created using Evo-viewer (https://temography.com/en/), which were used to quantify aspects of the CCV structure. Data was collected from at least 2 repeats. Reported quantitative measurement values in the main text are reported as mean ± SEM.

### High throughput pseudo 3D morphology screening of CCVs

SEM images of metal replicas of unroofed protoplasts, taken with a FE-SEM Merlin Compact VP equipped with an In-lens Duo detector, were obtained from *Johnson et al* (6). This included images of CCVs in Col-0 and *WDMX2* protoplasts incubated in control (4-hour incubation at room temperature) or endocytic membrane bending disruptive (4-hour incubation at 37°C) conditions, which were collected using the same magnification (∼45 k) and pixel size settings (∼2.4-2.6 nm). These protoplasts were prepared on glass coverslips as previously described (6).

For the automated segmentation of CCV, we first generated a training set consisting of 13 example image pairs (raw image and segmentation masks) of the CCVs in replica images of wild type and WDXM2 cells subjected to control or disruptive conditions. CCVs in the raw images were manually annotated using the Napari software (30). We then trained Cellpose (23) using input for 5000 epochs (see Protocol 1 for further details). The resulting trained model was then used to segment individual CCVs. To determine the localization accuracy of the automated CCV segmentation, we calculated the mean pixel-wise *Intersection over Union* (IoU) of manually annotated and predicted CCVs. Pixel-wise IoU is defined by the number of pixels in both, the *manual* and the *predicted* CCV segmentation mask, over the number of pixels of the union of both segments.

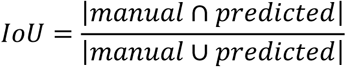

The predicted segmentation ROIs were then exported to Fiji (29) to quantify the area and average grey value of each CCV. Diameters of the equivalent circle were calculated from the CCV area. To estimate the curvature/pseudo 3D of the CCV, the average grey value of the CCV ROI was normalized by division to the average grey value of 4 manually selected PM ROIs in each image. The area and pseudo 3D values were plotted to define the overall morphology of a CCV. This was then divided in to 4 morphological categories (SR, small and round; SF, small and flat; LR, large and round; LF, large and flat) using an area threshold of 8500 nm^2^ (a CCV diameter of 105 nm) and a 3D value of 1.52 (the average of the 3 smallest CCVs in control conditions determined to be spherical by the experimenter).

## Supporting information

Supplemental Data and Protocols

## Data Availability

Example data and the code generated in this study is available at: https://doi.org/10.5281/zenodo.6563819

## Acknowledgements

This research was supported by the Scientific Service Units of Institute of Science and Technology Austria (ISTA) through resources provided by the Electron Microscopy Facility, Lab Support Facility and the Imaging and Optics Facility. A.J. is supported by funding from the Austrian Science Fund I3630B25 to J.F.

## Author Contributions

Conceptualization; A.J., W.A.K and J.F. Methodology; A.J. and W.A.K. Investigation; A.J., W.A.K. and T.C. Software; C.S. Validation and formal analysis; A.J. Resources; D.A.D. and S.Y.B. Writing original draft; A.J. All authors reviewed and edited the final manuscript.

