## Supplemental Data and Protocols for "User-friendly electron microscopy protocols for the visualization of biological macromolecular complexes in three dimensions: Visualization of *planta* clathrin-coated vesicles at ultrastructural resolution"

**Supplemental Information**

**Supplemental Movies**

Movie S1 – 3D reconstruction of a CCV in a protoplast

Movie S2 – 3D reconstruction of isolated CCVs from plant cells

**Supplmental Figures**

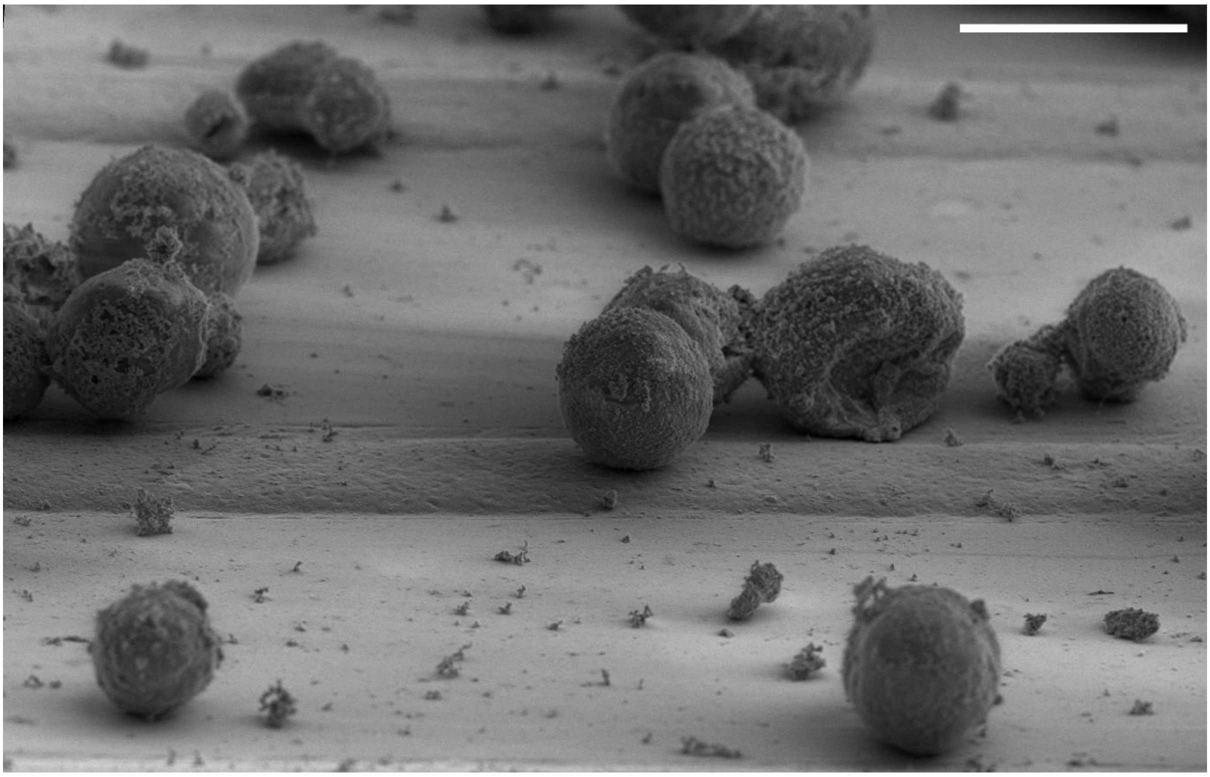

**Fig. S1. Protoplasts attached to an EM grid**

Representative SEM image of protoplasts derived from Col0 Arabidopsis seedling roots attached to EM grids and coated with platinum and carbon. The image was acquired using an Aquilos 2 Cryo-FIB SEM (Thermo Fisher). Scale bar, 20 µm

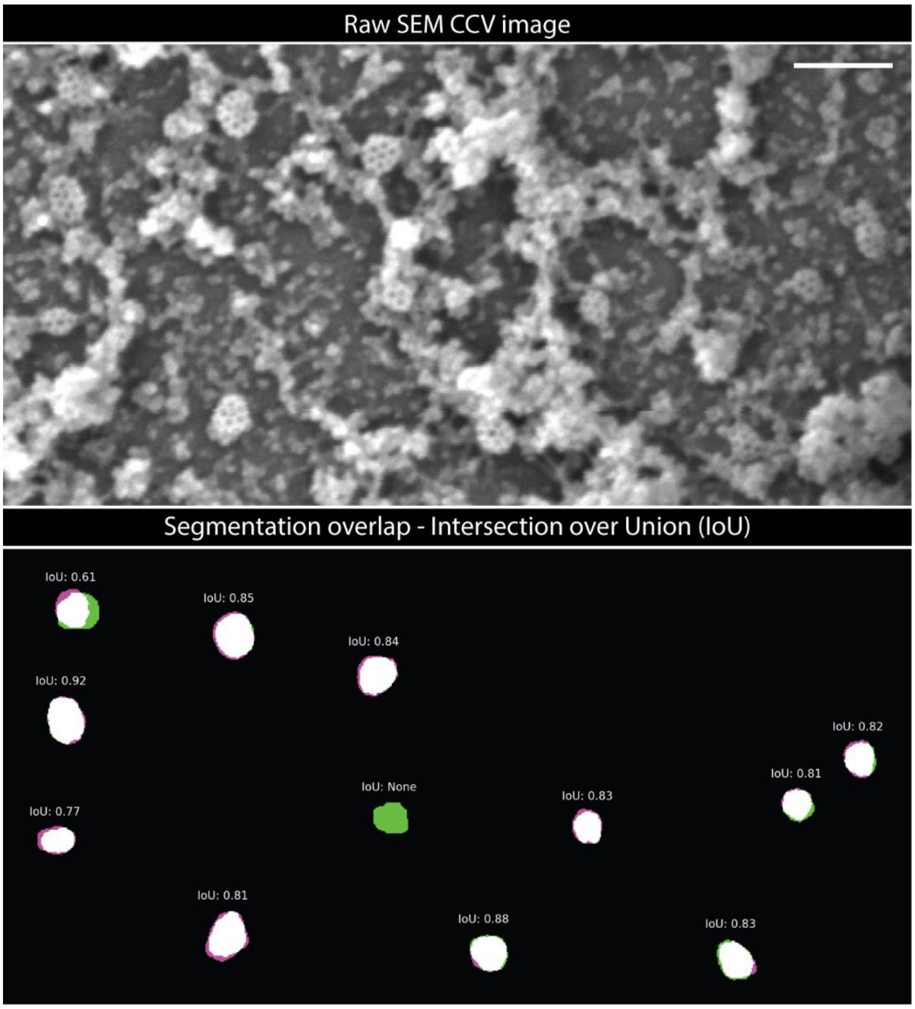

**Fig. S2. Pixel overlap of CCVs segmented by manual or automated means**

Representative metal replica of an unroofed protoplast cell derived from Arabidopsis seedlings, imaged with SEM (FE-SEM Merlin Compact VP equipped with an In-lens Duo detector). The segmentations from manual (green) and automated analysis (magenta) were overlayed, and an Intersection over Union (IoU) pixel overlap value was determined for each CCV. Scale bar, 200 nm.

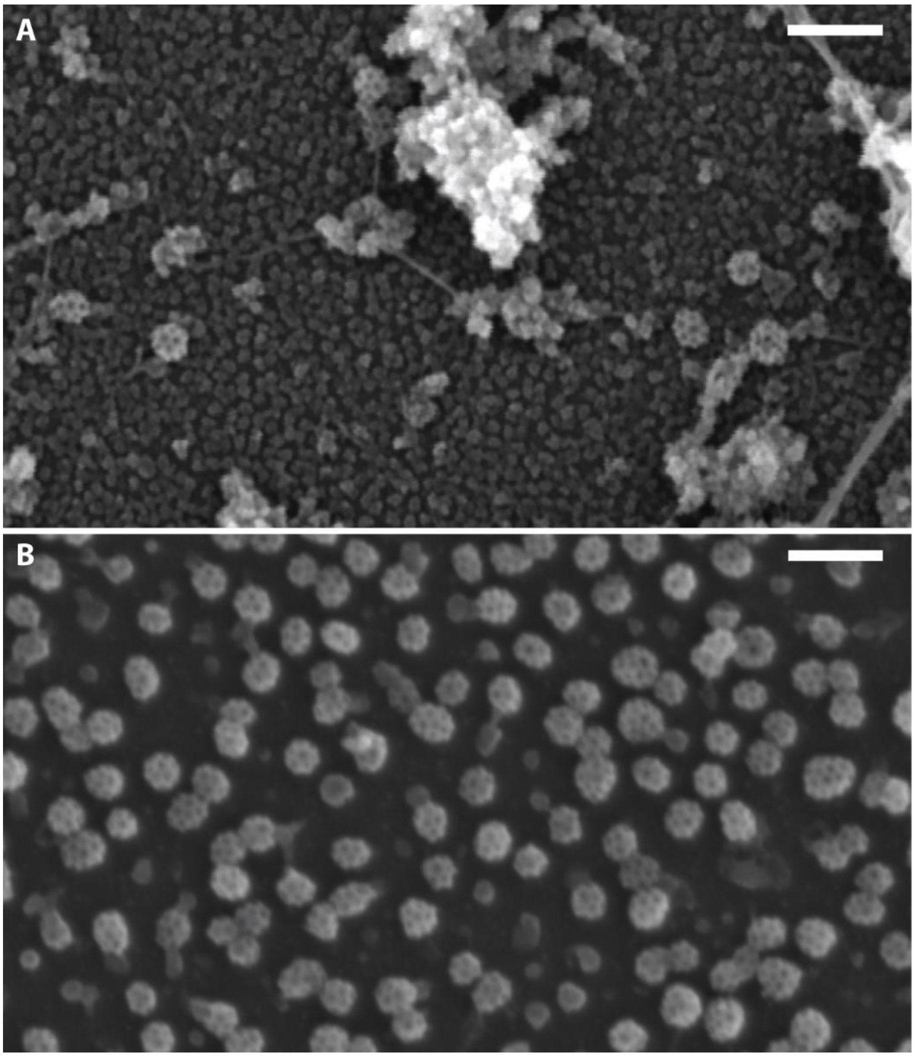

**Fig. S3. 'On-grid' preparations of CCVs from unroofed cells and isolated preparations**

(A) Representative SEM image of a metal replica of an unroofed protoplast derived from roots of Col-0 Arabidopsis seedlings. (B) Representative SEM image of a metal replica of isolated CCV preparations from Arabidopsis suspension cells. Scale bars, 200 nm.

**Supplemental Tables**

| | Size | | Visible Structure | | | | |
|---|---|---|---|---|---|---|---|
| **CCV** | **Diameter (nm)** | **Vol. ($nm^3$)** | **Pentagon** | **Hexagon** | **Not fully closed** | **Total** | **Mean arm length (nm)** |
| 1 | 66.90 | 1254081.88 | 5 | 5 | 8 | 18 | 16.64 |
| 2 | 71.85 | 1553842.88 | 4 | 4 | 8 | 16 | 17.00 |
| 3 | 71.69 | 1543323.78 | 4 | 2 | 8 | 14 | 16.65 |
| 4 | 79.21 | 2081704.89 | 4 | 4 | 8 | 16 | 16.69 |
| 5 | 84.65 | 2540718.45 | 5 | 6 | 6 | 17 | 17.72 |
| 6 | 77.94 | 1983739.44 | 7 | 5 | 5 | 17 | 17.37 |
| 7 | 79.71 | 2122016.34 | 5 | 5 | 10 | 20 | 16.11 |
| **Average** | 75.99 | 1868489.66 | 5 | 4 | 8 | 17 | 16.88 |
| **SEM** | 0.63 | 45538.34 | | | | | 0.06 |

**Table S1. Quantification of CCVs in unroofed protoplasts**

| | Size | | Visible Structure | | | | |
|---|---|---|---|---|---|---|---|
| **CCV** | **Diameter (nm)** | **Vol. ($nm^3$)** | **Pentagon** | **Hexagon** | **Not fully closed** | **Total** | **Mean arm length (nm)** |
| 1 | 63.75 | 1085210.4 | 7 | 5 | 6 | 18 | 17.53 |
| 2 | 78.94 | 2060762.9 | 8 | 4 | 9 | 21 | 18.31 |
| 3 | 62.91 | 1043171.6 | 7 | 2 | 7 | 16 | 17.91 |
| 4 | 78.27 | 2008426.8 | 4 | 3 | 7 | 14 | 19.18 |
| 5 | 61.95 | 996118.28 | 6 | 5 | 8 | 19 | 18.05 |
| 6 | 67.75 | 1302958.3 | 7 | 5 | 7 | 19 | 17.94 |
| 7 | 59.75 | 893497.27 | 7 | 2 | 6 | 15 | 17.79 |
| 8 | 73.86 | 1688070.1 | 7 | 4 | 6 | 17 | 18.22 |
| 9 | 86.94 | 2753269.9 | 10 | 6 | 10 | 26 | 18.46 |
| 10 | 65.45 | 1174691 | 5 | 3 | 6 | 14 | 18.3 |
| 11 | 68.68 | 1357115.5 | 5 | 3 | 6 | 14 | 18.6 |
| 12 | 71.14 | 1508041.3 | 7 | 1 | 7 | 15 | 18.22 |
| 13 | 70.62 | 1475650.1 | 6 | 3 | 8 | 17 | 18.66 |
| 14 | 70.00 | 1436664.2 | 4 | 4 | 5 | 13 | 16.91 |
| 15 | 80.02 | 2146829.4 | 6 | 5 | 7 | 18 | 18.83 |
| 16 | 77.15 | 1923880.3 | 5 | 7 | 7 | 19 | 17.32 |
| 17 | 68.22 | 1329999.3 | 5 | 2 | 7 | 14 | 18.79 |
| 18 | 73.59 | 1669727.4 | 9 | 6 | 6 | 21 | 19.36 |
| **Average** | *71.06* | *1547449.12* | *6* | *4* | *7* | *17* | *18.22* |
| **SEM** | *1.65* | *110644.50* | | | | | *0.14* |

**Table S2. Quantification of isolated CCVs**

| Col 0 | | | | | | | |
|---|---|---|---|---|---|---|---|
| | CCVs Detected | | | Average Area ($nm^2$) | | Average curvature | |
| **Image** | **Manual** | **Cellpose** | **Intersection over Union (IoU) (%)** | **Manual** | **Cellpose** | **Manual** | **Cellpose** |
| 1 | 15 | 16 | 81 | 3740.12 | 4035.60 | 2.03 | 2.00 |
| 2 | 22 | 20 | 83 | 4137.68 | 4538.13 | 2.00 | 1.92 |
| 3 | 23 | 19 | 83 | 4295.45 | 4559.88 | 2.02 | 1.98 |
| 4 | 9 | 9 | 83 | 5767.86 | 6236.06 | 2.30 | 2.27 |
| 5 | 9 | 9 | 88 | 4951.62 | 6236.06 | 2.11 | 2.27 |
| 6 | 5 | 4 | 85 | 5299.04 | 6063.83 | 2.41 | 2.38 |
| 7 | 8 | 10 | 80 | 4497.79 | 4641.10 | 2.50 | 2.33 |
| 8 | 6 | 6 | 91 | 4773.57 | 4624.17 | 1.93 | 1.92 |
| 9 | 6 | 6 | 81 | 7044.85 | 7295.16 | 1.93 | 1.92 |
| 10 | 10 | 9 | 81 | 6050.70 | 6590.24 | 2.92 | 2.86 |
| 11 | 15 | 13 | 87 | 6087.50 | 6492.98 | 2.55 | 2.58 |
| 12 | 16 | 18 | 83 | 4084.17 | 3808.60 | 2.10 | 2.12 |
| 13 | 18 | 18 | 82 | 4287.50 | 4399.86 | 2.07 | 2.01 |
| 14 | 37 | 39 | 84 | 4442.69 | 4383.13 | 2.04 | 2.00 |
| 15 | 10 | 10 | 82 | 4769.86 | 4967.41 | 2.48 | 2.25 |
| 16 | 13 | 14 | 85 | 4713.78 | 5090.21 | 1.99 | 2.00 |
| 17 | 12 | 12 | 76 | 4897.70 | 5187.19 | 2.48 | 2.47 |
| ***Total*** | *234* | *232* | | | | | |
| ***Mean*** | | | *83* | *4931.87* | *5244.09* | *2.23* | *2.19* |
| ***SEM*** | | | *0.8281* | *210.07* | *250.09* | *0.07* | *0.07* |
| ***Diameter*** | | | | *79.24* | *81.71* | | |

**Table S3. Manual and automated segmentation and analysis of wild type CCVs**

| *WDXM2* Control | | | | | | | |
|---|---|---|---|---|---|---|---|
| | CCVs Detected | | | Average Area ($nm^2$) | | Average curvature | |
| **Image** | **Manual** | **Cellpose** | **Intersection over Union (IoU)** | **Manual** | **Cellpose** | **Manual** | **Cellpose** |
| 1 | 7 | 7 | 79 | 4358.63 | 4846.07 | 2.50 | 2.13 |
| 2 | 10 | 11 | 81 | 5336.40 | 6069.10 | 2.15 | 2.14 |
| 3 | 7 | 7 | 79 | 5005.36 | 5879.29 | 1.95 | 1.92 |
| 4 | 13 | 13 | 81 | 5131.19 | 5685.69 | 2.14 | 2.10 |
| 5 | 9 | 9 | 82 | 4955.53 | 5622.99 | 1.68 | 1.66 |
| ***Total*** | *46* | *47* | | | | | |
| ***Mean*** | | | *80* | *4957.42* | *5620.63* | *2.09* | *1.99* |
| ***SEM*** | | | *0.6* | *163.505* | *208.831* | *0.134* | *0.091* |
| ***Diameter*** | | | | *79.44* | *84.59* | | |
| ***WDXM2*** **Induction to disrupt endocytic membrane bending** | | | | | | | |
| | CCVs Detected | | | Average Area ($nm^2$) | | Average curvature | |
| **Image** | **Manual** | **Cellpose** | **Intersection over Union (IoU)** | **Manual** | **Cellpose** | **Manual** | **Cellpose** |
| 1 | 15 | 18 | 70 | 8527.77 | 7320.09 | 1.19 | 1.20 |
| 2 | 32 | 26 | 72 | 4369.39 | 4909.21 | 1.28 | 1.27 |
| 3 | 24 | 26 | 78 | 6473.68 | 6726.43 | 1.42 | 1.45 |
| 4 | 35 | 35 | 80 | 6825.70 | 7098.20 | 1.38 | 1.35 |
| 5 | 24 | 26 | 73 | 7145.67 | 7842.16 | 1.33 | 1.34 |
| ***Total*** | *130* | *131* | | | | | |
| ***Mean*** | | | *74* | *6668.44* | *6779.22* | *1.32* | *1.32* |
| ***SEM*** | | | *1.9* | *672.056* | *501.193* | *0.040* | *0.041* |
| ***Diameter*** | | | | *92.14* | *92.90* | | |

**Table S4. Manual and automated segmentation and analysis of PM associated clathrin structures in WDMX2 cells which enables the inducible disruption of endocytic membrane bending**

**Supplemental Materials and Methods**

**Protocol 1 - STEM Tomography of unroofed protoplast cells derived from Arabidopsis seedlings**

Here we present a method that allows the tomographic imaging of single CCVs in plant cells derived from roots of Arabidopsis seedlings. Protoplasts are made from seedlings, attached directly to EM grids, unroofed and coated with metal to allow their observation via STEM. By acquiring tomographic tilt series, high resolution 3D reconstructed representations of single CCVs are generated can be quantified for various metrics, such as size and coat arrangement details.

Materials

½ AM Agar plates supplemented with 1% sucrose

10 cm Petri dish (VWR # 470210-568)

Analysis workstation (with Composer and Evo-viewer (https://temography.com/en/))

Arabidopsis seeds of interest

Benchtop 50 ml centrifuge (Thermo megafuge 40R, TX-1000 rota)

Benchtop vacuum chamber (7 mbar)

Cell culture dish polystyrene , 500 $cm^2$ square (Corning® VWR #734-1727)

Cell filter, pore size 40 µm (VWR #732-2757)

Curved tweezers (VWR # 63042-970)

EM gold grids 300 mesh with Formvar film (Electron Microscopy Sciences #FCF300-Au)

Filter paper Rotilabo® (Roth #AP60.1)

High vacuum metal coating device for platinum and carbon (Leica ACE600)

Laboratory oven with cooling function and integrated shaking platform (Memmert GmbH; IPP 55)

Orbital laboratory shaker (IKA; KS 130)

Parafilm (Biozym #743311)

Razor blade, disposable (Wilkinson Sword)

Scanning-transmission electron microscope (Jeol JEM2800 STEM)

Screw cap tube, volume 50 ml (Falcon® Sarstedt #62.547.255)

Syringe filter with pore size 0.22 µm (TPP #99722)

Task Wipes Kimtech® (Kimberly-Clark #7558)

Tweezers with clamping ring, Dumont stainless steel type L7 (Plano #T5129)

<u>Chemicals</u>

Agar (Duchefa #P1001.1000)

Cellulase R10 (Duchefa #C8001)

$D^+$ Saccharose (Sigma #84100)

Ethyleneglycol-*bis*(β-aminoethyl)-N,N,N′,N′-tetraacetic acid (EGTA; Merck #E3889)

Glutaraldehyde EM grade (Agar Scientific Ltd. #16220)

Hexamethyldisilane (Sigma #440191)

Macerozyme R10 (Serva #28302)

Mannitol (Merck #PHR1007)

Morpholinoethanesulphonic acid (MES; Duchefa #M1503.0100)

Murashige-Skoog (MS) powder (Duchefa #M0221.0050)

Murashige-Skoog (MS) powder with vitamins (Duchefa #M0222)

Phalloidin (Sigma #P2141)

Piperazinediethanesulfonic acid (PIPES; Sigma #P6757)

Polyvinyl Formal Resin (Formvar; Electron Microscopy Sciences #15800)

Polyethylene glycol MW 2,000 (PEG; Merck #821037)

Poly-l-lysine (Sigma #P8920)

Sucrose (Sigma #S0389)

Taxol (Merck #PHL89806)

Triton X-100 (Sigma #X100)

Tannic acid (Sigma #403040)

Uranyl acetate (AL-Labortechnik E.U. #77870.02)

Solutions

0.5 M MES

0.8 M Mannitol

AM agar (1 l stock: 10 g of D+ Saccharose, 2.3 g of Murashige-Skoog Powder, 0.5 g of MES and 8 g of Agar, pH 5.9 with KOH)

Enzyme solution (0.4 M Mannitol, 20 mM KCl, 20 mM MES pH 5.7, 1.5% Cellulase R10, 0.4% Macerozyme R10 in $H_20$)

Extraction buffer (2 µM phalloidin, 2 µM Taxol, 1% (w/v) Triton X-100 and 1% (w/v) polyethylene glycol (PEG; MW 2,000) in PEM buffer (100 mM PIPES free acid, 1 mM MgCl2, 1 mM EGTA; pH 6.9 adjusted with KOH)

Fixation buffer (2% (v/v) glutaraldehyde in 0.1 M phosphate buffer, pH 7.4)

GM Media (0.44% (w/v) Murashige-Skoog (MS) powder with vitamins, 89 mM Sucrose, 167 mM mannitol, pH 5.5 adjusted with KOH)

Graded ethanol solutions (10%, 20%, 40%, 60%, 80%, 96% in $ddH_20$)

PEM buffer (100 mM PIPES free acid, 1 mM $MgCl_2$, 1 mM EGTA; pH 6.9 adjusted with KOH)

Tannic acid 0.1% (w/v) in $ddH_20$

Uranyl acetate 0.2% (w/v) in $ddH_20$

W5 solution (154 mM NaCl, 125 mM $CaCl_2$, 5 mM KCL, 2mM MES)

Protocol

***Plant preparation (4 hours, 10-12 days before protoplasting)***

1 – The seeds of interest are sterilized by ethanol washing (10 mins 98% ethanol with gentle agitation) and dried in a sterile flow hood.

2 – The seeds are sprinkled onto ½ AM agar plates. To get a sufficient number of protoplasts, the seeds should be sown densely in two rows on at least 3 plates.

3 – The seeds are stratified by incubating the plates in the dark at 4°C for 2-3 days.

4 – The plates are then transferred to growth rooms (21°C, 16-hour light cycles), where they are incubated for 8-10 days.

***EM grid preparation (2 hours, the day before protoplasting)***

** *Critical point* –EM gold grids are very fragile, so handling of the grids requires great care to ensure that grids do not get bent or the film becomes broken.

5 – The formvar-coated EM gold grids are placed on filter paper, inserted into the ACE600 coating device and sputtered with 10 nm carbon (pulsing, working distance 90 mm, pressure 5.0E-5 mbar, rotating)

6 – The grids are then placed on to a 20 µl drop of poly-lysine on a parafilm layer and incubated for 15 min at room temperature (RT).

7 – The grids are washed by placing them on 20 µl drops of $ddH_2O$, dried for at least 1 h in the air and stored in dry glass petri dishes at RT until use.

***Making protoplasts from seedling roots (6 hours)***

8 – Fresh enzyme solution is prepared and filtered with a syringe filter of pore size 0.22 µm into a 10 cm petri dish.

9 – The seedlings are cut on their agar plates to separate the root from the rest of the plant.

10 – Several roots are transferred into the petri dish with enzyme solution using curved tweezers. While the roots are held with the tweezers, they are cut into small pieces (~1-2 mm) with a razor blade.

** *Critical point* – the cuts need to be as small as possible to facilitate the generation of protoplasts.

11 – The petri dish containing the cut roots and enzyme solution is incubated in a vacuum (7 mbar) for 20 min.

12 – The cuttings/enzyme solution is incubated for 3 h on a rocker in the dark with gentle agitation (~10 rpm).

13 – During this incubation, fresh W5 buffer and GM buffer are prepared.

14 – The cuttings/enzyme solution is filtered with a cell filter of pore size 40 µm in to a 50 ml Falcon® tube and supplemented with 20 ml of W5 buffer.

15 – The solution is centrifuged at 100 g for 2 min at RT.

*** Critical point* - A low centrifuge deceleration value (~half of max) is required to prevent damaging the protoplasts.

16 – The supernatant is removed (be sure to leave ~500 µl to avoid accidentally aspirating the cells) and the pelleted cells are resuspended in 20 ml W5 buffer.

17 – The solution is centrifuged at 100 g for 2 min at RT.

18 – The supernatant is discarded, and the cells are resuspended in 2 ml of W5 buffer and incubated for 30 min on ice.

19 – During this incubation, a humid chamber is prepared for further cells/grid incubations using polystyrene cell culture dishes lined with damp/wet paper tissues. Each EM grid is held with a tweezers closed with a clamping ring (to ensure the gird can be held horizontally without the experimenter) and placed inside the incubation chamber.

20 – The cells are centrifuged at 100 g for 2 min at RT and resuspended in 200 µl hyperosmotic GM solution (prepared in step 13).

21 – Plate 12 µl of the cell solutions on to each EM grid and incubate the grids/cells in the humid chamber prepared in step 19 for 3h at RT.

** *Critical point* – ensure the solution does not overflow onto the forceps, as this will result in the aspiration of the solution from the sample, consequently drying and damaging of the sample.

***Unroofing the cells (2 hours)***

** Note – for the following steps (22-33), the washing and treatments of the samples are conducted using droplets of the various solutions on parafilm sheets. 30 µl droplets were used for all solutions, except for the extraction buffer (step 23) – which used 20 µl.

22 – The GM media is blotted from each grid with filter paper, and samples are washed by placing the grids floating (sample side down) on droplets of PBS.

23 – Samples are incubated in extraction buffer for 5 min at RT on a rocker plate with gentle agitation.

24 – Samples are washed in PEM buffer supplemented with 1% (w/v) PEG 2000 3 times 2 min each at RT.

***Sample fixation (1 hour)***

25 – The cells are fixed in the fixation buffer for 30 min at RT in the humid chamber.

26 – Samples are washed with 0.1 M PB 2 times 5 min each at RT, and stored in 0.1 M PB at 4°C in the humid chamber until further use (typically overnight).

***Sample dehydration (4 hours)***

27 – Samples are washed in 0.1 M PB for 5 min at RT.

29 – Samples are washed in ddH$_2$O for 5 min at RT.

30 – Samples are incubated in freshly made 0.1% (w/v) aqueous tannic acid for 20 min at RT.

31 – Samples are washed in ddH$_2$O 3 times 5 min each at RT.

32 – Samples are incubated in 0.2% aqueous uranyl acetate for 20 min at RT.

33 – Samples are washed in ddH$_2$O 2 times 5 min each at RT.

34 – The grids are then transferred to glass bowls filled with ddH$_2$0.

35 – Samples are dehydrated by washing the grids in the bowls with graded ethanol (10%, 20%, 40%, 60%, 80%, 96% and 100%) for 2 min each at RT.

36 – Samples are incubated in pure hexamethyldisilane two times 2 min each at RT.

37 – Samples are then air dried by evaporation of the hexamethyldisilane.

***Metal coating (1 hour)***

38 – The samples are coated sequentially with 3 nm platinum and 4 nm carbon and stored in a dry humidity-controlled cabinet. Platinum coating: sputtering, working distance 90 mm, pressure 8.0E-3 mbar argon, tilt 0°, rotating; carbon coating: pulsing, working distance 90 mm, pressure 5.0E-5 mbar nitrogen, tilt 0°, rotating.

***Single CCV tomographic image acquisition (~1 hour per CCV tomogram)***

*** Note* – To save time, the sample grids can be screened using TEM and/or SEM at lower magnification to identify sample cells with many visible CCVs respectively purified vesicles before using STEM tomogram to focus on single CCVs.

39 – Sample grids are observed under a JEOL JEM2800 STEM at 200kV AV. The microscope needs to be calibrated and aligned properly following the manufacturer's instructions.

40 – Sample grids are cut in half with a razor blade and mounted on a half-mesh high-tilt holder (Jeol EM-21010/Z09291THTR).

41 – Once a CCV is located, tomograms (typical tilt angles -72 to +72° with step size 4°, along single tilt axis) are acquired using 'STEM Meister' (https://temography.com/en/), with auto focus and drift correction options activated.

** note - To allow CCVs to be compared directly and precisely, the same magnification setting should be used. In this study we used 600000x magnification with a pixel size of 0.84 nm.

***Reconstruction and Quantification (~60 minutes per tomogram)***

*** Note* – The tomograms were processed using Composer (https://temography.com/en/) and analyzed using Evo-viewer (https://temography.com/en/). It is also possible to use the IMOD as an open source alternative (https://bio3d.colorado.edu/imod/) (1).

42 - Each frame of the tomogram is manually screened to ensure the CCV of interest is in view and in focus.

43 - The tomogram is first aligned roughly by image cross-correlation using a coarse alignment algorithm, and then landmarks are added manually on visible structures to allow a precise alignment.

44 – Simultaneous iterations reconstruction technique (SIRT), with 20-40 iterations, is used to construct a 3D representation of CCVs.

45 – The clathrin arms are traced to create a wireframe model, and quantitative data is exported for further analysis.

**Protocol 2 – STEM Tomography of CCVs isolated from cultured Arabidopsis cells**

Here we present a protocol to visualize single CCVs in 3D in isolated CCV preparations from Arabidopsis cell culture systems. As the CCVs are processed to produce metal replicas in the same way as the unroofed cells in Protocol 1, it thus allows the direct comparison of purified CCV to those found in cells.

** Note - CCVs are isolated from undifferentiated T87 Arabidopsis suspension cell cultures using differential gradient centrifugation as described by (2). In the protocol described, the enriched CCV sample was at a concentration of 0.33 µg/µl in CIB buffer (100 mM MES, 0.5 mM MgCl, 3 mM EDTA, 1 mM EGTA, pH 6.4).

Materials (in addition to those required for protocol 1 [steps beyond 25])

Blotting paper (Whatman™ GE Healthcare #10311809)

EM copper grids 300-mesh with carbon film (Electron Microscopy Sciences #CF300-CU)

Glow discharge system ELMO™ (Agar Scientific Ltd.)

Protocol

***Mounting the CCVs on to EM grids (2 hours)***

1 – The EM grids are subjected to glow discharge for 4 min at pressure 7x10E$^{-1}$ mbar and current 30 mA in the ELMO$^{TM}$ glow discharge device.

2 – 5 µl of the purified CCVs are applied to the grids and allowed to absorb for 4 min at 4°C in a humid chamber.

3 – The excess solution is removed with blotting paper.

**Note – Take care that the samples do not fall dry during the whole procedure.

***Fixation, dehydrating, coating, image acquisition and tomographic reconstruction (8 hours)***

Sample grids are then prepared and imaged as described above, starting with step 25.

**Protocol 3 – High throughput pseudo 3D morphological screening of CCVs**

Here we present a robust protocol allowing the high throughput 3D morphology analysis of CCVs in 2D SEM images. As this protocol makes use of a single visual plane at a lower magnification than used in the STEM protocols, image acquisition of many CCVs can be conducted much more rapidly than with single CCV tomography approaches (Protocols 1 and 2). These images are then subjected to a deep learning analysis to automatically detect CCVs in an unbiased and high throughput manner. Using this automated detection, individual CCVs are quantified to define their morphology. Thus, this protocol allows the rapid screening of CCVs via size and a pseudo 3D measurement (curvature).

Example SEM images of CCVs in metal replicas of unroofed Arabidopsis root protoplasts derived from seedlings were obtained from (3). In contrast to Protocols 1, the protoplasts were prepared on glass coverslips as described in (4).

Materials

GPU enabled workstation computer with Napari (doi:10.5281/zenodo.3555620) with the Cellpose (5) plugin installed (see https://github.com/Mouseland/cellpose-napari for installation guide) and Fiji (6).

Example data and the code generated in this study is available at: https://doi.org/10.5281/zenodo.6563819

Protocol

***Generating a CCV prediction model (6 hours [5 hours of this is unsupervised training])***

1 – Example images containing CCVs are selected to train the prediction model.

*** Critical point* – the images used to train, and subsequently analyzed, should be acquired at the same resolution (i.e., pixel size and magnification should be similar).

2 – Image pairs are created from the example images selected in step 1. Here, one is the raw CCV image, and its ‘pair’ image is the same image with the CCVs manually annotated (Fig. 3A). This can be done in Napari, where masks are made by creating unique labels for each CCV. For each training image:

a. The raw image is loaded in Napari.
b. A new layer is added.
c. The paint button is selected (keyboard shortcut – ‘P’) and used to color in a single CCV.
d. A new label (‘+’ button) is selected (keyboard shortcut – ‘I’) and used to color in a different CCV.

*** Critical point* – each CCV must have a unique label.

e. Steps a-d are repeated until all CCVs in the image are labeled, and the masked layer is then exported as a ‘tif’ file.

*** Critical point* – the image pairs must have the same name, but with the CCV annotated mask images denoted with the addition of ‘_masks’ to the name (e.g. 01.tif and 01_ mask.tif).

3 – In the Napari environment, the following command is entered:

python -m cellpose --train --n_epochs 5000 --diameter 0 --use_gpu --dir ‘…\trainingData’[1] --pretrained_model None --chan 0 --mask_filter _masks

[1] This is the full directory path of where the image pairs are located.

This generated a model file which will be used for the following steps.

***Automated bulk Predictions (~10 seconds per image)***

** Critical point - Here we show how to analyze a group of unseen CCV images. However one can use the Naprai GUI to examine single images to check the quality of the prediction model before proceeding further.

4 – Raw images of CCVs to be analyzed are placed within a single folder.

5 – In the Napari environment, the following command is used:

python -m cellpose --use_gpu --dir “..\unseenData”[1] --pretrained_model "…\trainingData"[2] --save_png --no_npy

[1] This is the full directory path of where the unseen images (to be analyzed) are located.

[2] This is the full directory path of where the model generated in step 3 is located.

This generates outlines of the CCVs which are saved as text files.

***Analysis of CCV morphology (~2 minutes per image)***

6 – Fiji/ImageJ is prepared by installing the 'cellpose_outlines_to_roi.py' plugin to Fiji (this transforms the Cellpose outlines to ImageJ ROIs). The measurements (found in the analyze menu) are set to include area and mean grey value.

7 – A raw image is opened and the 'cellpose_outlines_to_roi.py' plugin is used to load the Cellpose CCV predictions.

8 – Measurements of the CCVs are made by pressing the 'measure' button in the ROI manager.

9 – The results are then exported to an excel file for further analysis and the Cellpose ROIs were cleared from the Fiji ROI manger.

10 – In the same raw image, 4 ROIs on areas of the PM (typically in each corner of the image and avoiding any large structures which are not PM) should be drawn. Their grey values are then measured and exported to Microsoft Excel. The mean value of these ROIs served as the background signal intensity to normalize the CCV grey values against, thereby allowing an estimation of their curvature to be calculated (Fig. 3C) (per image: mean CCV grey valve / mean grey value of the 4 PM ROIs).

11 – Steps 6-10 should be repeated until all the images are analyzed.

12 – The morphology of the CCVs can be analyzed by making a scatter plot of the mean area against the estimated curvature.
